# The biofilm matrix protects *Bacillus subtilis* against hydrogen peroxide

**DOI:** 10.1101/2025.01.12.632602

**Authors:** Erika Muratov, Julian Keilholz, Ákos Kovács, Ralf Moeller

**Affiliations:** German Aerospace Center (DLR e.V.), Institute of Aerospace Medicine, Radiation Biology Department, Aerospace Microbiology Research Group, Linder Hoehe, Cologne (Köln), Germany; University of Bonn, Institute for Microbiology and Biotechnology, Meckenheimer Allee 168, 53115 Bonn, Germany; Institute of Biology, Leiden University, Sylviusweg 72, 2333 BE Leiden, Netherlands

**Keywords:** *B. subtilis*, biofilm, hydrogen peroxide, endospores, exopolysaccharides, EPS, TasA, ROS, oxidative stress

## Abstract

Biofilms formed by *Bacillus subtilis* confer protection against environmental stressors through extracellular polysaccharides (EPS) and sporulation. This study investigates the roles of these biofilm components in resistance to hydrogen peroxide, a common reactive oxygen species source and disinfectant. Using wild-type and mutant strains deficient in EPS or sporulation, biofilm colonies were cultivated at various maturation stages and exposed to hydrogen peroxide. EPS-deficient biofilms exhibited reduced resilience, particularly in early stages, highlighting the structural and protective importance of the matrix. Mature biofilms demonstrated additional protective mechanisms, potentially involving TasA protein fibers. In contrast, sporulation showed limited contribution to hydrogen peroxide resistance, as survival was primarily matrix-dependent. These findings underscore the necessity of targeting EPS and other matrix components in anti-biofilm strategies, suggesting that hydrogen peroxide-based disinfection could be enhanced by combining it with complementary sporicidal treatments. This study advances our understanding of biofilm resilience, contributing to the development of more effective sterilization protocols.

## 1. Introduction

The ability of microorganisms to grow either as swarming cells or sessile biofilms on surfaces offers numerous benefits compared with planktonic growth in a liquid medium. Transitioning from the single cell motility through coordinated swarming to the immobilized lifestyle provides a flexible approach for nutrient utilization, a uniform proliferation, and increased resilience to environmental stressors (1–3). Thus, living in multicellular and multispecies communities is the most common form of microbial life, which is highlighted by their ubiquitous occurrence in medical, environmental and industrial settings (4, 5). The high impact of biofilms in these fields, notably within clinical environments, is underscored by statistics released by the National Institute of Health (NIH), revealing that biofilm-associated bacteria are responsible for 60% of all bacterial infections in humans. Furthermore, biofilms account for 80% of chronic and 65% of all nosocomial infections (6, 7). Besides the health aspect, biofilms have a high economic relevance. According to a comparative study from 2022, it is estimated that biofilms have an economic impact about USD 5,000 billion per year. The values refer to data from 2019, whereby the majority of the costs were caused by corrosive biofilms in industrial settings (8). These findings clearly illustrate the seriousness of biofilms as a health threat and the considerable challenge in inactivating them and developing anti-biofilm strategies. The increased virulence against eradication agents can be attributed to at least four categories (9):

I. Slow growth: Similar to the stationary phase in unicellular lifestyles, biofilms undergo physiological adaptations due to slower nutrient diffusion, resulting in reduced metabolic activity (4, 10). This adjusted growth kinetics produce more persistent cells with decreased susceptibility to sterilization regimes and antibiotics that target rapid cell growth (11, 12).
II. Communication: The capability to induce biofilm formation is highly dependent on an efficient cell-to-cell communication, termed quorum sensing (QS). QS systems differ between Gram-positive and negative bacteria and are based on the release of chemical signals (13). Once these signals are recognized, they can be utilized for optimal environmental adaptation. Hence, QS facilitates efficient nutrient utilization and storage, genetic material transfer, and division of labor. This division of labor triggers cell differentiation, encompassing motility, secondary metabolite synthesis, and production of protective biofilm matrix components (7, 14, 15).
III. Extracellular matrix (EM): A key factor in enhancing virulence in biofilms is the protective extracellular matrix produced by the inhabiting cells, which they produce autonomously and occupy (16). This complex matrix comprises biopolymers, such as extracellular polysaccharides (exopolysaccharides = EPS), proteins, lipids, and nucleic acids (17). The precise composition of the EM varies among species and is dependent on cultivation conditions, substrates and medium (4, 18). It is assumed that the EM is primarily responsible for the resistance to disinfectants and antibiotics, as it prevents penetration either by adsorption or by reacting with the polymers in the EM (19, 20).
IV. Unknown factors: In addition to I-III, there must be further protective factors. For example, EPS are crucial but not essential for biofilm formation and survival (21, 22). Identifying these protective factors could be challenging due to the dynamic nature of biofilms, which is influenced by various compounds and mechanisms.

To maintain a safe environment for immunocompromised patients which are more susceptible to chronic infections, effective interventions are necessary to minimize potential risks of infections (23–25). Such interventions include the disinfection of contaminated surfaces with hydrogen peroxide-based chemicals, a registered disinfectant with bactericidal, viricidal, sporicidal and fungicidal properties (24, 26). Hydrogen peroxide (H_2_O_2_) is classified as one of the reactive oxygen species (ROS), which can arise from intracellular or extracellular oxidizing events, such as radiation exposure or mitochondrial phosphorylation (27). H_2_O_2_ is a robust oxidizing agent that in the presence of Fe^2+^ generates highly reactive hydroxyl radicals (^·^OH) which are able to damage macromolecules, such as DNA, lipids of the cell membrane, and proteins (27–29). The imbalance between ROS and protective endogenous compartments results in oxidative stress which subsequently cause cell death (30).

*Bacillus subtilis,* a Gram-positive facultative anaerobic soil bacterium, forms complex biofilm consortia and exhibits remarkable resistance owing to its ability of sporulation (31). Thus, this species is commonly utilized as a biological indicator in decontamination studies (32). In addition to endospore (hereafter referred to as spores) formation, the multicellular lifestyle offers numerous potential protective properties. The EM primarily consists of exopolysaccharides (EPS) and the protein TasA, which forms amyloid fibers essential for the biofilm scaffold. Moreover, *B. subtilis* biofilms produce a hydrophobin protein coat, formed by BslA, crucial for overall protection against desiccation and selective permeability (33–35). The organization of this biofilm assembly relies on nutrient availability and extracellular signals (36–38). Upon signal recognition and adequate environmental conditions, part of the population start to express biofilm-related genes leading to phenotypic heterogeneity that allows coexistence of motile and matrix-producing cells, as well as development of highly resistant spores at the later stage that are assumed to contribute to dispersal (39–41). This division of labor is tightly regulated and dynamic, with gene expression profiles adapting to environmental conditions (4, 42). These properties demonstrate the remarkable adaptation of biofilms to environmental stressors, making them challenging to inactivate once formed. *B. subtilis* biofilms have been utilized to dissect the influence of various treatment strategies, including disinfection agents, nanoparticles, and laser irradiation (43–47).

This study seeks to elucidate the impact of hydrogen peroxide on bacterial biofilms lacking EPS and spores, thus contributing to the development of targeted strategies for biofilm control and disinfection. Here, we used the architecturally complex colonies of *B. subtilis* to evaluate the role of these protective structures.

## 2. Material and Methods

### Spore production and purification

For spore production, 200 µL of an overnight culture were inoculated onto solidified Schaeffer sporulation medium (SSM) (48). The strains used in this study are listed in **Table *1***. Plates were incubated for 5-7 days at 37°C to achieve optimal spore quality and quantity. Spores were harvested from the plate using an inoculation loop and resuspended in 40 mL _dd_H_2_O containing sterile glass beads with a size of 3 mm in diameter. This facilitated resuspension using vortexing (2 minutes) and aided in dispersing cell debris released from the lysed mother cells. To achieve high spore quality and purity, the suspension was repeatedly washed until purity of >99% spores was confirmed using phase contrast microscopy. The pure spore solution was then stored in glass tubes at 4°C until utilized.

**Table 1:** Isogenic *B. subtilis* strains tested in survival ability to hydrogen peroxide. Tet^R^-tetracycline resistance, Cat^R^-chloramphenicol resistance.

| Strain | Genotype | Deficiency | Reference |
| --- | --- | --- | --- |
| NCIB3610 | Wild type | None | (37) |
| ZK3660 | $\Delta epsA$-<br>O::Tet<sup>R</sup> | No production of<br>exopolysaccharides within the EM.<br>Remaining matrix components are<br>accomplished by TasA and BslA. | (49) |
| F-030 | $comI^{Q12L}$<br>$sigG$::Cat<sup>R</sup> | Deficiency in sporulation (inhibition<br>of late forespore polymerase<br>activities) | (50) |

### Bacterial biofilm cultivation

To obtain biofilms which are standardized and reproducible, a cultivation method according to Fuchs et al. was conducted (51). Briefly, an inoculum of spores with 10^8^ Colony Forming Units (CFU) mL^-1^ was utilized and pipetted in the middle of a hydrophilized PTFE filter (polytetrafluoroethylene, Merck Millipore®, pore size 0.4 µM, Merck KGaA, Darmstadt, Germany). This PTFE filter separates the growing biofilm physically from the medium, while enabling water and nutrient diffusion. The inoculated filter material was air dried under sterile conditions for 10 minutes and placed on solidified minimal medium (MSgg), adapted after Branda et al. (37, 52). As *B. subtilis* biofilms are highly heterogenous populations with changing cell and EM profile over time, differently matured biofilms were tested here, ranging from 24h to 72h. For 0h, pure inoculum of spores was pipetted onto the filter material. Here, the treatment was performed directly after the drying process. For the sporulation deficient strain of *B. subtilis* (Δ*sigG*) an overnight culture with planktonic cells from stationary phase with 10^8^ CFU mL^-1^ was used as inoculum and 0h control.

### Sample treatment and CFU determination

Biofilms were grown to distinct development stages and exposed to H_2_O_2_. PTFE filters carrying the biofilms were placed into sterile six-well plates. ROS stress was induced by adding 910 µL of 3% H_2_O_2_, diluted in PBS, to each biofilm. After treatment durations of 0, 10, 20, 40, and 60 minutes, the reaction was stopped by adding 10 mg mL^-1^ catalase solution. The treated biofilms were transferred into 2 mL reaction tubes containing glass beads (3 mm diameter). To ensure a reliable yield of viable cells, the reaction mixture suspension was also transferred to the 2 mL tube containing the biofilm. Each tube was vortexed for 2 minutes and survivability was assessed by calculating total CFU and the number of spores. To quantify the spores count, an aliquot of the sample was treated at 80°C for 10 minutes to inactivate vegetative cells.

### Statistical analysis

The CFU of the total cell mass and spores within the biofilm were calculated at each treatment point of the stress assays as well as the untreated control. Therefore, a dilution series was prepared and plated on LB agar. The average CFU was determined while all data are presented as the average of three biological replicates (n=3) with according standard deviations. The statistical analysis has been performed by using Tukey’s test with SigmaPlot (version 14.5) and OriginLab (version 2023).

## 3. Results

Hydrogen peroxide is known to be an efficient antimicrobial agent and is commercially used as a disinfectant. Numerous studies have demonstrated its efficacy in combating biofilms formed by *Pseudomonas aeruginosa* and *Staphylococcus aureus*, targeting both the matrix and the cells (24, 53, 54). In this study, our objective was to assess the anti-biofilm activity of hydrogen peroxide against *Bacillus subtilis* biofilms and the contribution of EPS and spores on survivability. Hence, macrocolony biofilms were cultivated at various growth stages, exposed to hydrogen peroxide and quantified via CFU determination. The cell count within wild-type biofilms consistently rises as maturity progresses, while total CFU comprises a mixture of vegetative cells and spores (**Figure 1**). The inoculum (0h) includes around 10^5^ spores per ml, resulting in little difference in cell count between total CFU and spores at this time point. Spore count peaks in mature (72h) biofilms, contributing to a notably reduced ratio of vegetative cells. The morphology of wt biofilms varies also with age. In the 24h stage, the characteristic concentric rings develop. Matured biofilms exhibit increased wrinkling and size. Overall, particularly at 48 and 72 hours, biofilms appear as highly heterogenous 3-dimensional structures.

**Figure 1:**
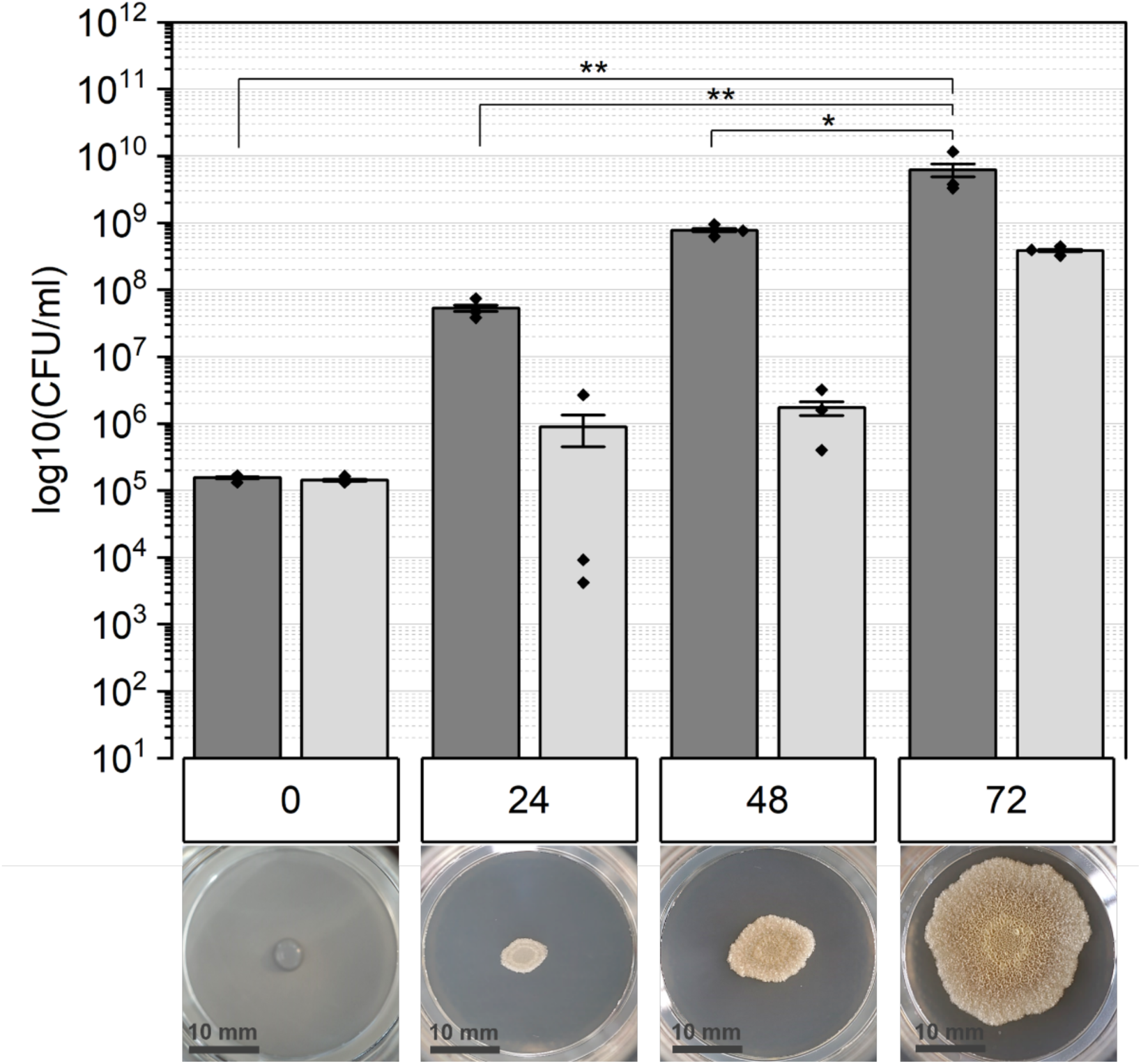
The quantity of untreated wild-type (wt) biofilms is depicted according to each stage of biofilm maturation (in hours). The biofilms are composed of a mixture of vegetative cells and spores, termed as “CFU total” (dark grey bars). Additionally, the proportion of spores was quantified and is represented as light grey bars. The macroscopic morphology is illustrated below for each respective time point.

*Bacillus subtilis* biofilms lacking extracellular polysaccharides within the matrix demonstrate variations in cell count when compared to the wt. Initially, at the 0-hour timepoint, CFU total and spores exhibit similarity, approximately 10^4^ CFU ml^-1^. However, after 24 hours, this pattern reverses, with spore count lower than the inoculum and maintaining consistency over time. Meanwhile, the number of vegetative cells experiences a remarkable increase, approximately fivefold higher than the spore count. As the biofilms mature, at 48 and 72 hours, the total cell count slightly surpasses that of the young biofilm stage and significantly exceeds the CFU count observed at the 0-hour stage. The lack of EPS leads to a noticeably altered biofilm morphology, as illustrated in **Figure 2**. At the 24-hour mark, there are notably fewer concentric rings, while the size remains largely similar to that of the wt. In 48-hour-old biofilms, the characteristic wrinkles typical for the wild type at this stage are absent. Nevertheless, the size remains comparable to that of the wild type. By the 72-hour timepoint, the biofilm is much smaller and exhibits a more uniform structure without wrinkles, showing only one visible concentric ring. Overall, biofilms lacking eps appear less dense and thinner compared to wt.

**Figure 2:**
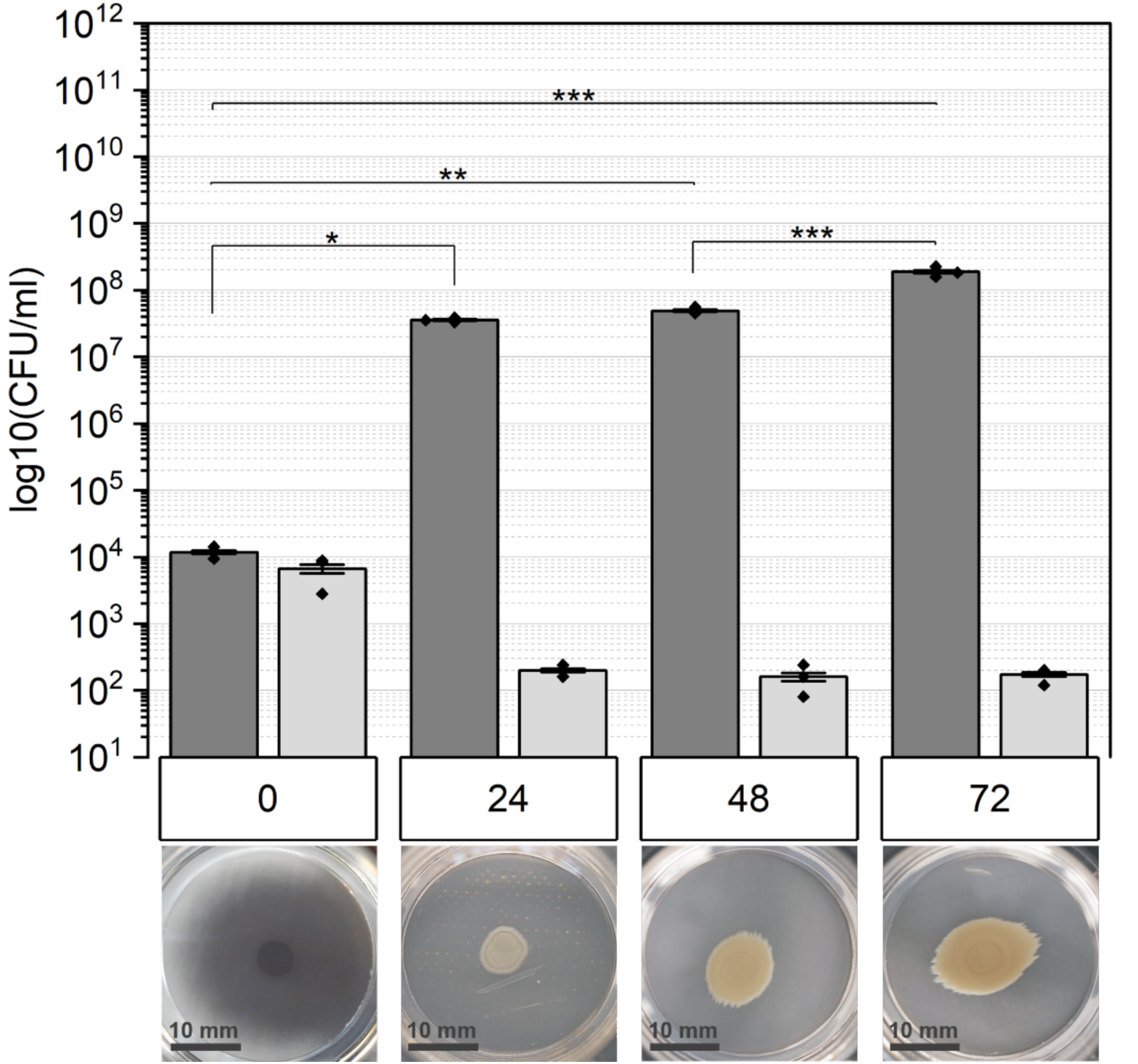
The number of untreated biofilms lacking exopolysaccharides (Δ*epsA-O*) is illustrated for each stage of biofilm maturation (in hours). Dark gray bars represent the total colony-forming units (CFU), while light gray bars indicate the spore count. Statistical significance was determined using Tukey’s test.

**Figure 3:**
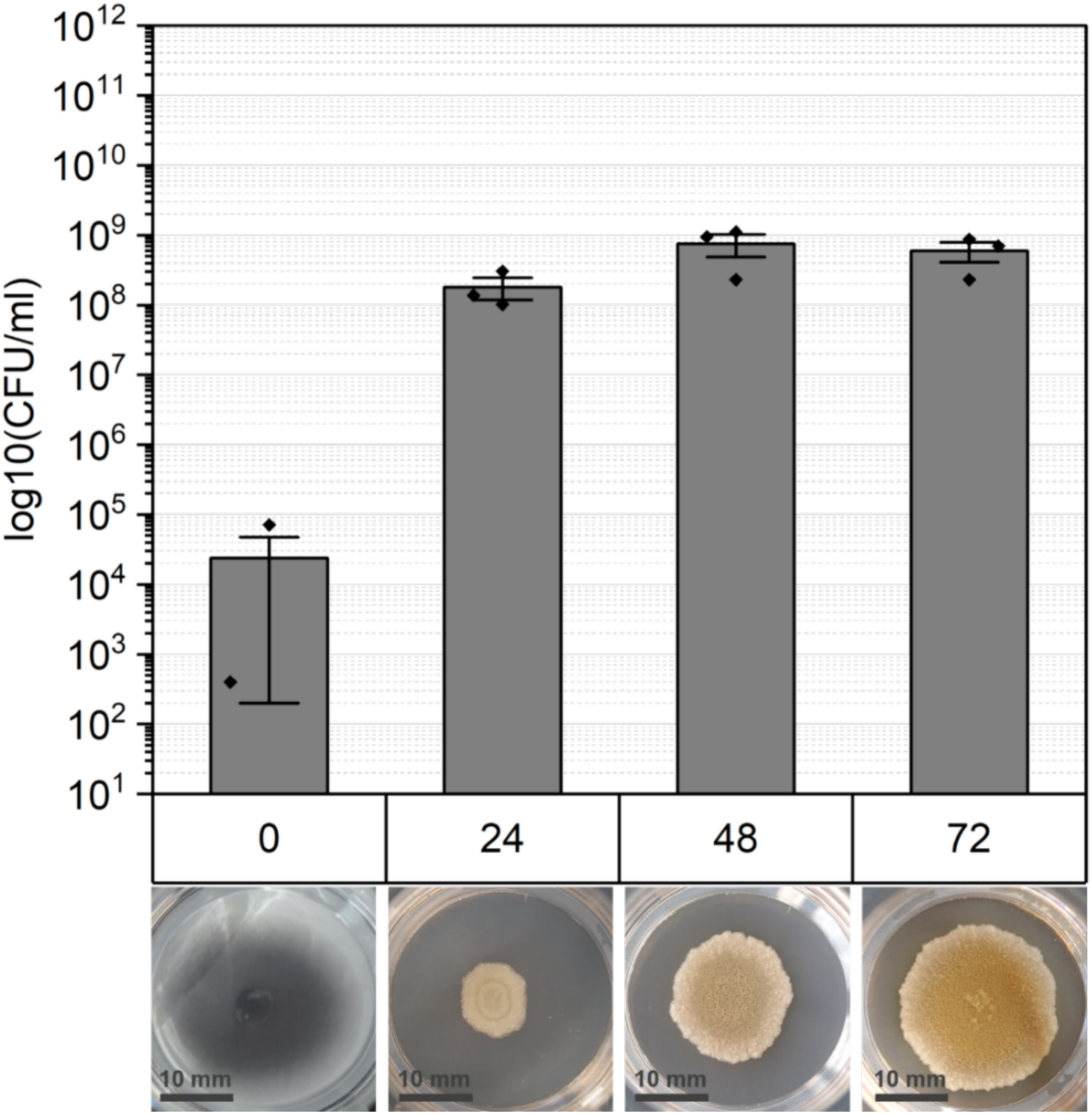
The quantity of untreated biofilms lacking spores (Δ*sigG*) is depicted for each stage of biofilm maturation (in hours). The bars represent only vegetative cells and is termed as “CFU total”. Macroscopic variation among biofilm age is shown below.

*B. subtilis* cell aggregates devoid of spores can be attributed to a deletion in the gene encoding the sigma factor G (**Table 1**, (50). Starting from 0 hours, with approximately ∼10^4^ planktonic cells per ml, the biofilm expands to double this amount at mature levels. Once a certain threshold is reached, the cell count stabilizes with minimal further increase.

Wt biofilms treated with hydrogen peroxide were revived, and their survival was assessed via CFU determination (**Figure *4***). The 0h timepoint (**Figure 4A**) indicates the initial spore count and represents the inoculum reference. Regardless the incubation time, the CFU remains stable, with nearly identical quantities observed between spores and CFU total. Remarkably, even after a 60-minute exposure, spore survival remained unaffected. 24-hour old consortia showed slight impact in survival but a significant decrease among CFU total (**Figure 4B**).

**Figure 4:**
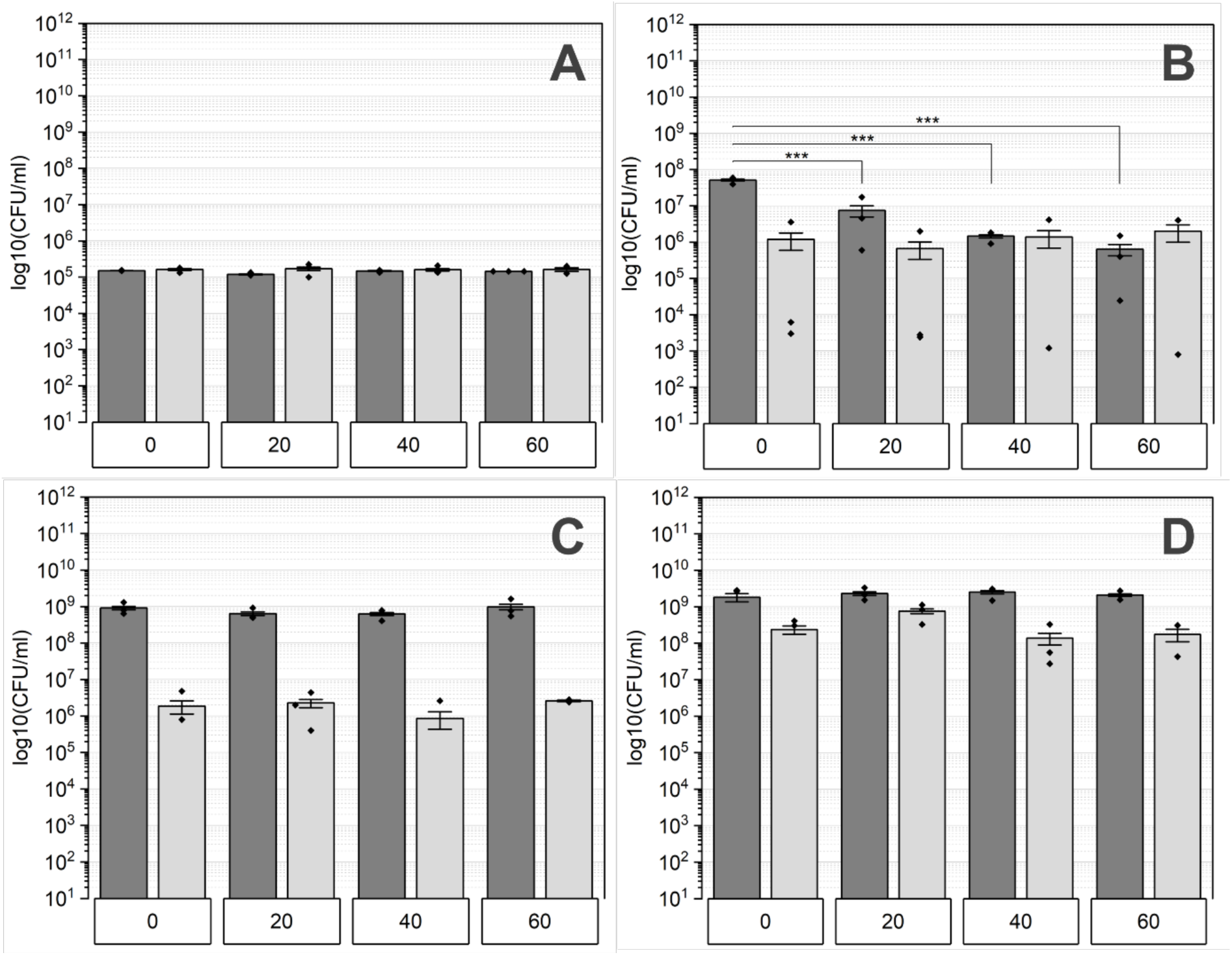
The quantity of wt biofilms treated with hydrogen peroxide is analyzed based on their maturation levels, represented as A: 0h, B: 24h, C: 48h, and D: 72h. Exposure durations to H_2_O_2_ include 0, 20, 40, and 60 minutes. Total colony-forming units (CFU) are represented by dark grey bars, while the count of spores is illustrated by light grey bars. Statistical significances were determined utilizing Tukey’s test.

However, the spore count remained nearly the same, while for 40 and 60 minutes treated biofilms, the spore quantity was similar to those of CFU total. Mature wt biofilms, cultivated for 48 and 72 hours, exhibited efficient resistance to hydrogen peroxide treatment, as the cell quantity was unchanged throughout incubation time. The only distinction between these maturation stages lies in the higher spore count observed in the 72h biofilms compared to those cultivated for 48 hours.

The control inoculum of the *eps* deficient strain shows similar outcomes to those of the WT strain at 0h and maintains a consistent cell count regardless of the exposure time (Figure 5A). Following biofilm formation, a mixture of vegetative cells and spores is present, with the spore count in this strain considerably lower than that in the WT strain, as observed in untreated samples. In early-stage biofilms (24h, Figure 5B), the total CFU is roughly three times higher than the spore count at the 0-minute mark.

**Figure 5:**
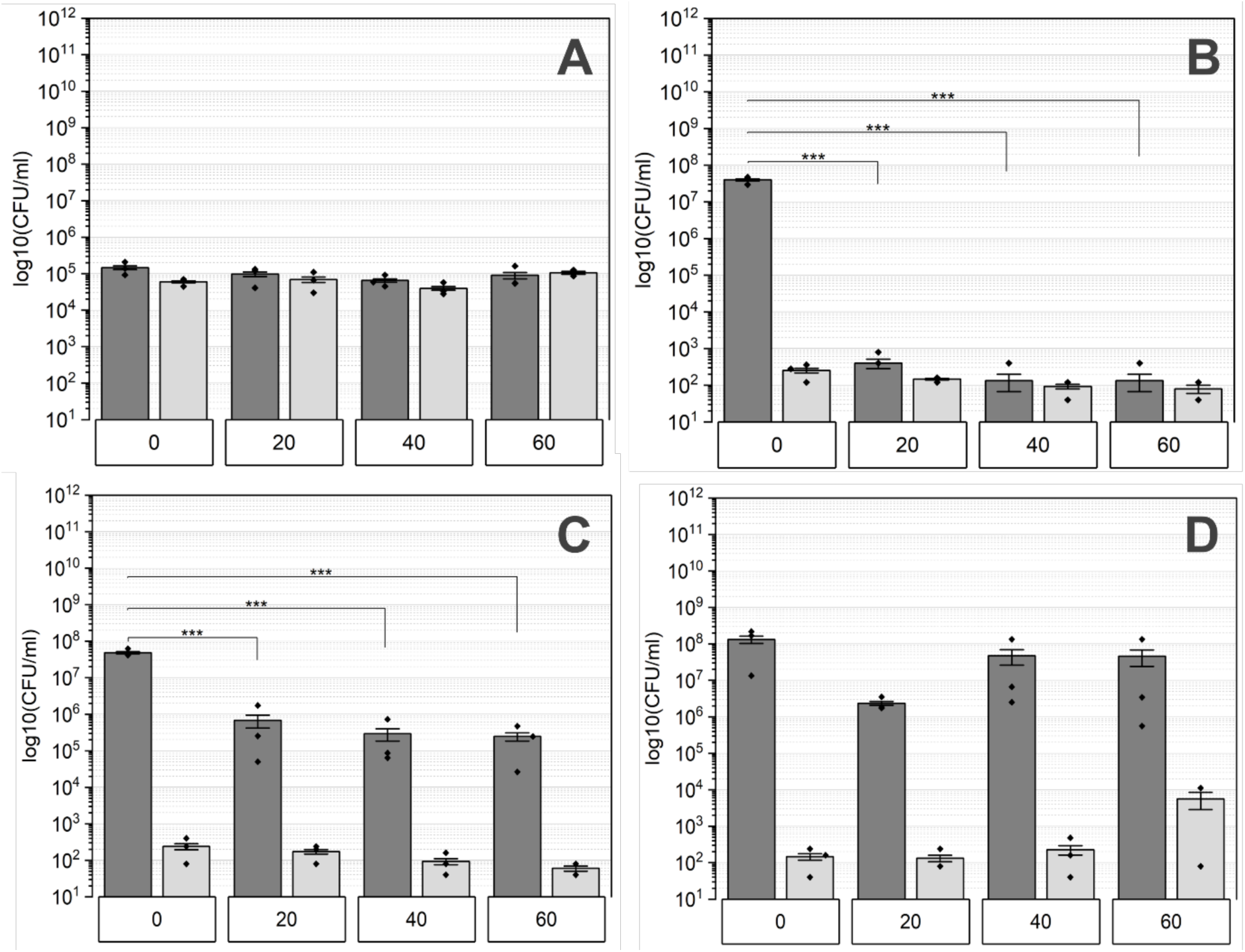
The *eps* deficient biofilms’ quantity treated with hydrogen peroxide was assessed at various maturation stages labeled as A: 0h, B: 24h, C: 48h, and D: 72h. These biofilms underwent exposure to H_2_O_2_ for durations of 0, 20, 40, and 60 minutes. Total CFU is depicted by dark gray bars, while the number of spores is shown by light gray bars. Statistical significance was evaluated through Tukey’s test.

However, for exposure durations between 20 and 60 minutes, the total CFU significantly declines, reaching levels comparable to those of spores.

The subsequent maturation stage, 48-hour-old biofilms show a higher survival rate compared to those aged 24 hours (Figure 5C). Nevertheless, a significant reduction in total CFU persists in comparison to the 0-minute control. Additionally, there is a slight decrease in the spore count following a 60-minute exposure to hydrogen peroxide. Biofilms grown for 72 hours show increased susceptibility to hydrogen peroxide after a 20-minute treatment compared to those exposed for 40 and 60 minutes (Figure 5D). Moreover, the total CFU count slightly increases with longer treatment durations, reaching its peak spore count at the 60-minute mark.

The final strain tested for resistance to hydrogen peroxide lacked SigG, making it incapable of producing spores (Figure 6). Planktonic cells from the stationary phase were used as inoculum (0h, Figure 6A) to evaluate the survival ability by determining the total CFU. After 20 minutes of treatment, no CFU could be detected and this remained consistent for longer incubation periods. Interestingly, once consortia are formed, the cells showed an improved resilience against hydrogen peroxide. Biofilms aged from 24 to 72 hours, showed similar results and were only slightly affected (Figure 6B-D). Exposure for 20 minutes resulted in minimal reduction in CFU, which was constant for longer incubation time.

**Figure 6:**
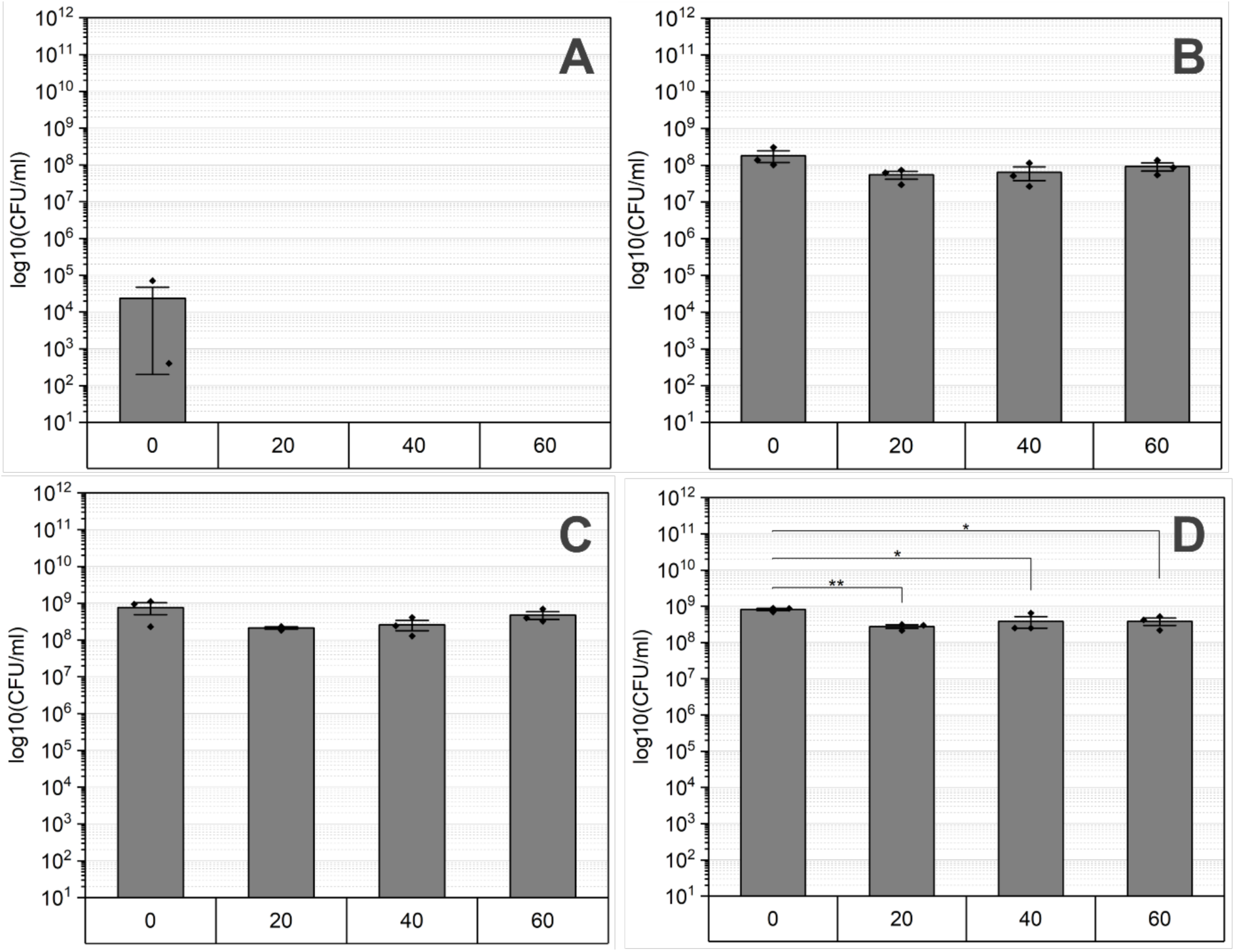
Biofilms deficient in SigG were exposed to hydrogen peroxide and tested in survival across different maturation stages labelled as A: 0h, B: 24h, C: 48h, and D: 72h. Exposure durations to H_2_O_2_ ranged from 0 to 60 minutes. Statistical significance was analyzed using Tukey‘s test.

## 4. Discussion

Hydrogen peroxide serves as a widely used commercial disinfectant, capable of targeting a broad spectrum of microbes, including spores. Some studies even report about its effectiveness against biofilms. However, there is a lack of available data regarding *Bacillus subtilis* biofilms and the contributions of EPS and spores to resistance. Moreover, many studies overlook the impact of varying biofilm ages, which could be pivotal in understanding resistance mechanisms. Before the survival assay was conducted, the biofilms in maturation stages ranging from 0 to 72 hours were cultivated and analyzed in their morphological phenotype as well as cell and spore quantity.

### Comparison of the morphology and cell/spore quantity in wildtype biofilms versus EPS- and spore-lacking variants

The formation of architecturally complex structures as biofilms is attributed to a spatiotemporal cycle involving alternating phases of motile swarming and sessile matrix production (55, 56). This process results in the formation of concentric rings, as observed in wt biofilms, while being less abundant in *eps-*deficient and Δ*sigG* colony biofilms (**Figure1-3,** (57)). Overall, biofilms lacking EPS show observable differences in texture and size. These consortia are impacted in their physical integrity and display a more homogenous appearance compared to wt. Although there are other matrix structures besides EPS, no wrinkles are visible. This effect strongly suggests that the formation of wrinkles is dependent on all matrix structures (58). The vertical expansion, or the consolidation phase of biofilms is facilitated by matrix-producing cells, which are disrupted in the *epsA-O*-deficient strain, leading to the thin colony morphology (55). Nevertheless, the preserved size in early-stage biofilms is maintained by swarming cells, which support two-dimensional expansion, or migration phase. Thus, the overexpression of the motile cell phenotype could compensate for the absence of EPS, thereby aiding in the growth of biofilms (57). Mature biofilms lacking EPS appear to be smaller than wt biofilms in this growth stage. Due to the importance of EPS for the structure, quorum sensing mechanisms and the adhesion, the expansion may occur up to a certain threshold and being halted (59). When comparing the sporulation-deficient biofilms to wt, they appear particularly in the early stage larger but lack concentric rings in the mature (48h and 72h) phases. However, in 72h wt biofilms, wave fronts are more prevalent in the center of the biofilm, whereas in Δ*sigG* consortia they appear denser at the edges (Figure 1 vs. Figure 3). The spatial arrangement of *B. subtilis* biofilms at specific time intervals prompts cell differentiation, including sporulation, at different stages and locations, culminating in characteristic colony morphology (60). Sporulation seems to be associated with the development of complex architectural formations, as indicated by the diverse biofilm structure observed in our investigation. Furthermore, Aguilar et al. validated the correlation between sporulation and matrix production by the protein KinD (61). Interestingly, Vlamakis et al. discovered that biofilms lacking spores due to *sigF* deletion do not undergo changes in biofilm structure but reduced spore quantity when matrix production is shut down (41, 49).

In our study biofilms which lack EPS in the matrix show as well reduced levels of spores compared to wt (Figure 2). The cell differentiation within biofilms is a highly regulated process, with matrix production and sporulation being connected through the activity of the bifunctional protein KinD. KinD mediates the phosphorylation (or dephosphorylation) of the master transcription factor Spo0A. As low quantities of phosphorylated Spo0A induces expression of matrix genes, EPS-deficient mutants exhibit delayed sporulation, leading to low spore count (61). In addition, the matrix mutant biofilms show less quantity of cells and spores, indicating higher number of lysed cells, thus increased cannibalism within the population (62, 63).

Comparing the overall cell numbers between the investigated strains, wt biofilms tend to an exponential increase in cell number, while the cell quantity of mutant strains reaches a plateau in mature stages. The EPS- and sporulation-deficient strain may differ in nutrient requirements which result in altered biofilm growth and CFU amount (64). One potential aspect for altered growth patterns observed in genetically manipulated biofilms could be limited nutrient storage and transport. In biofilm lifestyles, one defense mechanism against starvation involves the storage of compounds like glucose and calcium ions, which impact spatial metal ion distribution (58). Calcium is typically bound by the extracellular matrix (EM), while zinc, manganese, and iron are more freely distributed through water channels, accumulating in biofilm wrinkles (58, 65). Due to missing EPS this distribution is disordered, leading to restricted resilience, since these compounds are important for withstanding shear forces and toxic chemicals (58). In addition to the storage and transport, EPS contributes to quorum sensing (QS), which regulates the cell density and expansion of biofilms. The polysaccharides facilitate the stability of signal molecules necessary for QS, enhancing biofilm functionality and maintenance (66). Furthermore, QS is crucial for the division of labor within biofilms, which include the differentiation to sporulating cells. Spores are known to be a metabolically inactive (or reduced) dormant stage *Bacilli* (64, 67). The altered nutrient requirements could be explained by the low abundance of spores and missing EPS within the mutant biofilms, as there is known to be a link between EPS production and nutrient availability (64, 68).

### Contribution of extracellular polysaccharides to survival to hydrogen peroxide

The resistance of cells in a biofilm to a variety of disinfectants and antibiotics is attributed to the protective EM (69). The matrix of *B. subtilis* biofilms is mainly composed of polysaccharides and proteins, while former is more abundant in the mobile section (18, 70). Because of the multiple functions of EPS within the matrix, its protective capability against hydrogen peroxide was tested. *Eps* mutants are more susceptible to hydrogen peroxide at all maturation stages compared to wt (**Figure 5B-D**). Only spores (0h) demonstrated a similar resistance and were not affected in terms of survival (Figure 5A). Interestingly, the resistance characteristics differ across the maturation stages tested. In young biofilms (24h) only spores survived the hydrogen peroxide exposure, whereas mature biofilms did not exhibit the same resistance. Treated 48h and 72h biofilms show a higher number of total CFU than CFU of spores, indicating the survival of vegetative cells. Thus, apart from EPS, additional components within the matrix must contribute to the protection. The structural integrity of biofilms is crucial for surviving harsh environmental stressors, such as reactive oxygen species induced by hydrogen peroxide. Although the exact composition of the matrix varies depending on numerous factors and differs even among species, EPS and proteins are crucial for this integrity and are therefore highly abundant (18, 71). Besides the structural functionality, these compounds enhance the resistance to biocides. On one hand, the matrix acts as physical barrier against antimicrobial agents like hydrogen peroxide. On the other hand, they can react with them resulting in their depolymerization and thus disruption of aggressive hydroxy radicals. In addition to polysaccharides, amyloid fibers formed by the TasA protein are known to contribute to the virulence (72). Because of its characteristic beta-sheet structure, the interaction with antimicrobial agents that could lead to proteolysis is hindered, thereby TasA provides protection (72–75). Thus, a potential factor contributing to the survival of EPS-deficient (particularly mature) biofilms could be the presence of TasA fibers. Branda et al. describes TasA and EPS as the most important and abundant structures in the biofilm matrix (36). Further testing of a TasA mutant is necessary to shed more light on the role of amyloid fibers in hydrogen peroxide resistance. Overall, EPS are crucial for surviving oxidative stress, although they play a minor role in mature biofilms, as indicated by the slight reduction in cell quantity of biofilms. It is likely that TasA also contributes to this protection. Further analysis is required to confirm this hypothesis including exposing Δ*tasA* strain and a double mutant (Δ*epsA-O tasA*) to H_2_O_2_ to investigate co-dependency.

### Role of sporulation in hydrogen peroxide resistance

Cell differentiation within *Bacillus subtilis* biofilms is a crucial mechanism for adapting to dynamic environmental changes and stressors. This differentiation includes the formation of endospores, which allows the cells to persist under harsh conditions in a metabolically inactive (or reduced) state (76, 77). The transition into this dormant state is triggered by nutrient depletion and is accomplished by a range of different resistance mechanisms (78, 79). This is confirmed by the spore counts shown in Figure 1 as biofilms mature and nutrient levels decrease. The survival of oxidative stress induced by hydrogen peroxide is ensured by enzymes such as catalases or superoxide dismutase localized in the spore coat (80). For instance, spore-specific catalases like KatX are crucial for surviving hydrogen peroxide exposure during spore germination (81). Additionally, we observed macroscopic differences in biofilm morphology appeared between sporulation-deficient populations and wt biofilms (Figure 1 versus Figure 3). This observation indicates that the differentiation into spores could contribute to the structural integrity and thus, to biofilm resilience against hydrogen peroxide (82, 83). Hu et al. investigated the resistance of spores and vegetative cells from biofilms of *Clostridium perfringens* to oxidative stress and confirmed that spores were more resistant than vegetative cells and the sessile lifestyle has an enhanced resilience (84). However, numerous studies report the efficacy of hydrogen peroxide as sporicidal agent. Indeed, Sawale et al. determined D-values ranging from 0.08 to 0.95 minutes by using concentrations from 22 to 33%. In our study, wt spores were not reduced after 60 minutes treatment, but the used concentration was more than ten times lower (Figure 4A). Using a similar concentration of H_2_O_2_, show a decreased susceptibly and achieve “hardly any inactivation” in spores which is confirmed by further studies (85, 86). Either increasing the incubation time or concentration of hydrogen peroxide could improve the sporicidal efficacy. Looking on multicellular lifestyle it was expected that based on the spore-specific protection mechanisms, spores would contribute to overall resistance to hydrogen peroxide. Interestingly, our results in turn, showed that the formation of spores had no impact on the survival rate of biofilms (Figure 6). The quantity of cells was regardless the maturation or incubation time not affected in survival which emphasizes the importance of an intact and functional biofilm matrix.

## 5. Conclusion

This study has shown that EPS in the matrix play a major role in the protection against hydrogen peroxide whereas sporulation does not. A functional structural integrity with intact EPS are even more protective than the ability of forming spores in surviving oxidative stress. Furthermore, our results have shown that besides the EPS, especially in mature biofilms, additional protective structures remain which are most likely given by the TasA fibers. This research has revealed that EPS are probably the most protective component within the matrix and need to be tackled. Newly developed sterilization approaches are often based on hydrogen peroxide and should be combined with additional sporicidal agents like UV or heat.

## Declaration of competing interests

The authors declare that there is no conflict of interests

## Data availability

All data are depicted in the graphs or available on request.

## Acknowledgement

This paper is dedicated to the memory of Ralf Moeller. The authors express their appreciation to Prof. Dr. Jörg Stülke (University of Göttingen, Germany) and Prof. Dr. Madeleine Opitz (Ludwig-Maximilians-University München, Germany) for providing strains. Furthermore, we would like to thank Andrea Schröder for her help and support of this work.

This project was supported by the Deutsche Forschungsgemeinschaft (DFG) (MO 2023/3-1), and the German Aerospace Center (DLR), through Program Space - Research under Space Conditions, Partial Program 475, Project ISS LIFE 2.0 - Life in space: ISS and beyond to the Moon and Mars.

## CRediT authorship contribution statement

**Erika Muratov**: Writing – original draft, Visualization, Conceptualization, Investigation, Methodology, Data Curation, Formal analysis. **Julian Keilholz**: Investigation, Methodology, Data Curation, Formal analysis. **Ákos Kovacs**: Writing - Review & Editing. **Ralf Moeller**: Funding acquisition, Project administration, Conceptualization, Writing - Review & Editing, Supervision

## Notes

### Competing Interest Statement

The authors have declared no competing interest.

